# Development and application of eDNA-based tools for the conservation of white-clawed crayfish

**DOI:** 10.1101/732941

**Authors:** Christopher R. Troth, Alfred Burian, Quentin Mauvisseau, Mark Bulling, Jen Nightingale, Christophe Mauvisseau, Michael J. Sweet

## Abstract

1. eDNA-based methods represent non-invasive and cost-effective approaches for species monitoring and their application as a conservation tool has rapidly increased within the last decade. Currently, they are primarily used to determine the presence/absence of invasive, endangered or commercially important species, but they also hold potential to contribute to an improved understanding of the ecological interactions that drive species distribution. However, this next step of eDNA-based applications requires a thorough method development.
2. We developed an eDNA assay for the white-clawed crayfish (*Austropotamobius pallipes*), a flagship species of conservation in the UK. Multiple subsequent *in-situ* and *ex-situ* validation tests aimed at improving method performance allowed us to apply eDNA-based surveys to evaluate interactions between white-clawed crayfish, crayfish plague and invasive signal crayfish.
3. The assay performed well in terms of specificity (no detection of non-target DNA) and sensitivity, which was higher than more established traditional species survey methods. Quantification of species biomass was, however, less reliable.
4. Comparison of eDNA sampling methods (precipitation vs. various filtration approaches) revealed that optimal sampling method differed across environments and might depend on inhibitor concentrations.
5. Finally, we applied our methodology together with established assays for crayfish plague and the invasive signal crayfish and demonstrated their significant interactions in a UK river system.
6. Our analysis highlights the importance of thorough methodological development of eDNA-based assays. Only a critical evaluation of methodological strengths and weaknesses will allow us to capitalise on the full potential of eDNA-based methods and use them as decision support tools in environmental monitoring and conservation practices.

## Introduction

Since its initial conception as a method for aquatic ecological surveys (Ficetola et al., 2008), the use of environmental DNA (eDNA) based methods are rapidly increasing in popularity (Biggs et al., 2015; Jerde et al., 2013;. Advantages such as higher cost effectiveness compared to established traditional species survey methods and the non-invasive sampling approach have been emphasised (Goldberg et al., 2016; Huver et al., 2015). Nevertheless, the true potential of eDNA-based methods is just starting to be realized. Currently, eDNA-based tools are mostly used for simple presence/absence surveys, while they could also be used to study ecological interactions that determine species distribution and the conservation status of target species. However, such advances in application require careful method evaluations and the improvement of sampling approaches to increase reliability of detection and prevent false conclusions.

In the case of species-specific eDNA assays, the design and validation of the assay represents a critical first step (Geerts et al., 2018). During assay design, it is fundamental to ensure a high target specificity (Bylemans et al., 2018) by selecting suitable amplicon lengths, *in-silico* simulations and testing against amplification of non-target DNA. *In-vitro* laboratory validation should then ascertain that the assay complies with established guidelines (Bustin et al., 2009) and that limits of detection (LOD) and quantification (LOQ) are established. Further, field comparisons with established traditional survey methods are recommended to complement reliability assessments (Smart et al., 2015). However, both traditional survey approaches and eDNA-based methods are affected by various error sources potentially creating inconsistencies that require careful interpretation (Hinlo et al., 2017a).

Further, the reliability of eDNA-based tools is strongly influenced by sampling methodology (Hinlo et al., 2017b). Currently, precipitation and various filtration methods are applied to concentrate eDNA during field sampling. Filtration approaches have the advantage of collating eDNA from larger volumes of water compared to precipitation-based methods (Mächler et al., 2016). However, they can also incorporate the risk of missing particles below the filter pore size (Minamoto et al., 2016) and may lead to higher concentrations of inhibitors preventing targeted eDNA amplification (Mauvisseau et al., 2019a). Previous method comparisons have come to contrasting recommendations for difference species (Deiner *et al*., 2015; Dickie *et al*., 2018). Additionally, even for the same species the ‘optimal’ method for collecting eDNA may vary between lentic (i.e. ponds or lakes) and lotic (i.e. rivers and canals) systems (Geerts et al., 2018) and therefore careful method comparisons are recommended (Deiner et al., 2015).

In this study, we target the white-clawed crayfish, *Austropotamobius pallipes* (Lereboullet, 1858), an endangered and important umbrella species in the UK and Western Europe (Füreder et al., 2010). Range reduction of white-clawed crayfish began in the 1860s, with declines rapidly accelerating in the UK after the introduction of invasive non-native signal crayfish (*Pacifastacus leniusculus*, Dana, 1852) from North America in the 1970s (Holdich et al., 2009). Moreover, the spread of crayfish plague *Aphanomyces astaci* (Schikora 1906), an oomycete pathogen carried by signal crayfish, has greatly exacerbated the negative impact of invasive competitors, pollution and habitat degradation (Holdich et al., 2009). Despite its legislative protection (EU Habitats Directive), white-clawed crayfish has continued to decline by as much as 50-80% between 2000 and 2010 (Füreder et al., 2010). Due (at least in part) to the now rarity of the native species, traditional survey methods are having unsatisfactory success in monitoring populations (Gladman et al., 2010; Holdich and Reeve, 1991), highlighting the urgent need to develop new survey tools.

Consequently, the aim of this study was to develop a highly reliable eDNA assay for the detection of white-clawed crayfish, that allows the assessment of interactions with competing species and pathogens which threaten their survival. We designed a primer and probe set for the amplification of white-clawed crayfish DNA and critically evaluated the sensitivity and specificity of the assay through extensive *in-silico, in-vitro* and *in-situ* tests. Moreover, we evaluated the impact of different sampling methodologies on the reliability of the assay in mesocosm experiments and field tests implemented in different habitat types. Finally, this allowed us to assess in a UK river system the relationship between white-clawed crayfish, signal crayfish and crayfish plague, demonstrating the applicability of eDNA-based approaches for in-depth ecological investigations and ecosystem management.

## Materials and Methods

### Primer design and in-silico tests

Primer/probe design and validation followed guidelines established by MacDonald and Sarre (2017) aimed for assay development of species-specific eDNA methods. The primers and probe, targeting the Cytochrome C Oxidase Subunit 1 (COI) mitochondrial gene of white-clawed crayfish, were designed *in-silico* using Geneious Pro R10 (Kearse et al., 2012). The forward primer WC2302F 5’ - GCTGGGATAGTAGGGACTTCTTT - 3’, reverse primer WC2302R 5’– CATGGGCGGTAACCACTAC - 3’ and probe WC2302P 5’ - 6-FAM-CTGCCCGGCTGCCCTAATTC-BHQ-1-3’ amplified a 109bp fragment. To ensure specificity, *in-silico* tests were run against published sequences of closely related and/or co-occurring crayfish species.

### In-vitro validation

The specificity of the assay was further tested *in-vitro* against extracted DNA of either taxonomically similar, or co-occuring crayfish species. These included; *Faxonius limosus* (Rafinesque, 1817), *P. leniusculus*, *Astacus astacus* (Linnaeus, 1758), *Astacus leptodactylus* (Eschscholtz, 1823), *Procambarus clarkii* (Girard, 1852), and *Procambarus virginalis* (Lyko, 2017). DNA was extracted from crayfish tissues using the Qiagen DNeasy^®^ Blood & Tissue kit, following manufacturers’ instructions. PCRs were performed using the primers and methods from Folmer et al. (1994) and sequenced by Eurofins Genomics (Germany) to confirm species identity of all specimens. Specificity of the newly designed assay was then assessed using qPCR.

The reactions for both tissue and all eDNA samples contained; 12.5μL TaqMan^®^ Environmental Master Mix 2.0 (Life Technologies^®^), 6.5μL DH_2_0, 1μL (10μm) of each primer, 1μL (2.5μm) of probe with the addition of 3μL template DNA. qPCR’s were performed with 6 technical replicates (i.e. qPCR replicates) of each sample on a StepOnePlus™ Real-Time PCR System (Applied Biosystems). qPCR conditions were as follows: 50°C for 5 min, denaturation at 95°C for 8 min, followed by 50 cycles of 95°C for 30 s and 55°C for 1 min. Six no template controls (NTC’s) were prepared using RT-PCR Grade Water (Ambion™) alongside a duplicated serial dilution of control white-clawed crayfish DNA (10^-1^-10^-3^ ng uL^-1^) for each qPCR plate that was run. In each subsequent experiment, negative PCR controls were included in this fashion.

### Limits of detection (LOD) and quantification (LOQ)

The reliability of our assay was also assessed, following the Minimum Information for Publication of Quantitative Real-Time PCR Experiments (MIQE) Guidelines, which recommend the establishment of a calibration curve to determine LOD and LOQ (Bustin et al., 2009). We prepared a serial dilution of DNA extracted from white-clawed crayfish starting from 0.79ng μL^-1^ to 7.9×10^-8^ ng μL^-1^ with 10 qPCR replicates per dilution analysed. The LOD was defined as the last standard dilution that resulted in a detection of target DNA with at least one qPCR replicate at a threshold cycle (Ct) of <45. The LOQ was defined as the last standard dilution in which targeted DNA was detected and quantified in a minimum of 90% of qPCR replicates of the calibration curve under a Ct of 45 (Mauvisseau et al., 2019b).

### In-situ validation

The reliability of the assay was further field tested by comparing eDNA-based and traditional capture-mark-recapture sampling techniques at six sites of confirmed white-clawed crayfish presence (2017) in the Centre-Val de Loire region, France. Each site was visited at least twice in subsequent nights between 22^nd^ June and 1^st^ of August 2018 (see supplementary information, Table S1 Individual white-clawed crayfish were surveyed using a torching approach, counted and marked using a white waterproof marker stain. In the second night the survey was repeated and marked, and non-marked crayfish were differentiated. Population size was estimated using the Lincoln-Petersen Method 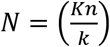, where N is the estimated population size, n is the number of crayfish marked on the 1^st^ visit, K is the total number captured on the 2^nd^ visit and k is the number of those captured individual which were already marked on the 2^nd^ visit. (Petersen 1896, Lincoln 1930). Additionally, eDNA samples (two environmental replicates) were collected at each site using the 0.22μm Sterivex filters (see below for detailed description). eDNA samples were collected between the 22^nd^ and 29^th^ June 2018. The water volume filtered varied due to cases of high turbidity (consistent minimum volume of 150mL, see Table S2 for list of all sample volumes). Furthermore, additionally to the method below, eDNA filters were fixed with 2mL of ethanol to accommodate for the longer storage and transport time between the field and the laboratory. All sampled locations are part of an extensive monitoring programme for white-clawed crayfish population studies and due to conservation reasons, locations of sites are not reported. The water temperature, environmental variables and the volume of sample filtered for eDNA samples (varied due to variable turbidity) were recorded at each site.

### Comparison of eDNA sampling methodologies in mesocosms

Further, our aim was to assess the impact of eDNA sampling methodology on both the probability of eDNA detection and the signal strength (i.e. Ct) of its detection. We tested differences between the most common eDNA sampling methods utilised to date, including (i) ethanol precipitation (Biggs et al., 2015), (ii) 2μm pump-based filtration (Strand et al., 2014), (iii) 0.45μm pressure filtration and (iv) 0.22μm pressure filtration (Spens et al., 2017). All methods were assessed in two mesocosms, housed at Bristol Zoological Gardens, Bristol, UK during autumn 2018. Both mesocosms were designed to the same specifications but contained different water volumes and crayfish numbers. Mesocosm 1 had a volume of 3000L and contained 249 individual adult white-clawed crayfish and sub-adults (between 17 months and four years old) with a total biomass of 1.3kg. Water parameters of mesocosm 1 were; pH 8, temperature 11°C, and under natural light conditions. Mesocosm 2 contained a volume of 1000L of water with the same pH (8) but with higher water temperatures (16°C) and under artificial light conditions. This mesocosm held a larger number of crayfish (379) but all were juvenile (five months old). The sensitivity of juveniles to handling did not allow us to obtain the exact biomass of this mesocosm, but biomass was estimated as 250g. Both mesocosms were set up as recirculating ‘flow through filtration systems’, ensuring high water quality at all times. Six samples for each method and mesocosm combination were collected from both mesocosms.

In brief; eDNA samples classified hereafter as ‘precipitation’ samples were collected following the protocol outlined in Biggs et al. (2014). 1L of water (20 x 50mL subsamples) was collected from ~20cm below the surface and after homogenization, a subsample of 90 mL (6x 15mL) was aliquoted into sterile tubes containing a pre-mixed buffer solution (Biggs et al., 2014). Samples were stored at −20°C prior to extraction and extracted following Tréguier et al. (2014). For batch of samples analysed, negative extraction controls were included and processed analogously to other samples.

eDNA samples taken with a 2μm pump-based filtration consisted of 2L of water collected by the same sub-sample method outlined above but were then filtered through a Millipore Glass fibre filter AP25, 47mm (2μm pore size) using a peristaltic pump (Masterflex E/S Portable Sampler, Cole-Parmer, USA). The filter was housed in an In-Line Filter Holder 47mm (Merck) connected by silicone tubing. The combined use of a peristaltic pump and a larger filter pore size allowed us to substantially increase the amount of water filtered. The filter was then removed from the pump system and stored at −20°C before extraction. Equipment was soaked and cleaned with 10% bleach between samples. Filters were extracted following Spens et al. (2017). eDNA sample collections for 0.22μm and 0.45μm pressure filtration were undertaken in the same manner. 20 sub-samples were collected and a 50mL syringe (BD Plastipak™, Ireland) was then used to pressure filter 250mL of water through a sterile enclosed filter (Sterivex™, Merck^®^, Germany) with either a pore size of 0.22μm (Polyethersulfone membrane) or 0.45μm (Polyvinylidene fluoride membrane). All filters were stored at −20°C, and extracted following Spens et al. (2017).

### In-situ comparison of eDNA sampling methodologies

Complementary to the tests in the mesocosm experiment, we also evaluated sampling methodologies under natural conditions. However, we performed only pairwise method comparisons in order to contain sampling effort in the field. As a test in a lentic system, eDNA samples were collected from a 1000m^2^ pond in the South West of England after the release of 40 white-clawed crayfish individuals (equal juvenile-adult and male-female ratios, total biomass of 436g). Here, precipitation (sample volume: 90mL) was compared against 0.22μm pressure filtration (sample volume: 250mL). Sampling started on the 20^th^ April 2018 and was repeated two hours, seven days, 14 days and 35 days after crayfish release. At each sampling time, three field replicates were taken from four sites around the pond for each method. Additionally, 20 50mL sub-samples taken from the entire pond perimeter were pooled, homogenised and sampled with three field replicates per method.

Our second field test was conducted in a lotic system. We sampled 10 sites (situated approx. 1km apart) along a chalk stream river in Dorset (UK), during September 2017, and 4 sites along a river in Derbyshire (UK). Here, ethanol precipitation was used in comparison to pump-based filtration (2μm, sample volume: 2L), using three field replicates at each site per method (*n* = 42). Samples collected in the river system (20 pooled sub-samples as described above) were taken in an interval of 1-2m along a diagonal downstream-to-upstream transect across the river. In this field test, we also assessed the ability to screen for crayfish plague using both sampling methods. qPCRs in this instance were run using the primers and probe developed by Strand et al. (2014). For all of the field method comparisons, bottled distilled water was sampled on site and processed as field negative controls.

### Field test of white-clawed crayfish, signal crayfish and crayfish plague co-existence

Finally, we assessed the distribution of white-clawed crayfish, signal crayfish and crayfish plague in a river-system in Derbyshire (UK). Two field replicates were taken at each of eight sites along the river in November 2017. Six of these sites were located in proximity to the inflow of tributaries and two field replicates were taken upstream and downstream of their confluence to capture the influence of populations potentially present within tributaries (supplementary information, Fig. S1). The other two sites were sampled with two field replicates. Sampling was conducted using the precipitation method outlined above and water samples were tested for the occurrence of all three species. Protocols of Mauvisseau *et al*., (2018) for signal crayfish and of Strand *et al*., (2014) for crayfish plague were applied.

### Statistical Analysis

Samples measured for the establishment of a standard curve were analysed using a linear regression to evaluate the relationship between DNA dilution and Ct. A log-log data transformation decreased the models Akaike Information Criterion (AIC; reduction by 23 units compared to second best model) and was therefore used for downstream analyses. Residuals were tested for autocorrelation, normal distribution and any remaining patterns (same procedure applied in all regression analyses). A logistic regression analysis was also applied to test the relationship between DNA concentration and binomial detection data assessing the change of detection probability with DNA concentrations. For the mesocosm and field samples, the relationship between (i) the population density established by traditional sampling methods and (ii) the Ct values and detection probability (calculated as the fraction of technical replicates that resulted in positive detection) of eDNA measurements were examined in a linear regression model. Differences in sample volumes between locations (due to turbidity) were accounted for by including sample volume as a predictor in regression models, and log-log and untransformed models were compared using AIC. Further, Ct and detection probability of different sampling methods were compared using ANOVA analyses followed by Tukey’s HSD post-hoc tests, and t-tests or nested ANOVA’s (lotic and lentic systems, where only two methods were compared). Prior to ANOVAs, heteroscedasticity was evaluated, and data transformed if necessary. Finally, the co-existence of white-clawed crayfish with signal crayfish and crayfish plague was tested in regression models using detection probability of all three species. All described statistical analyses were performed using R version 3.4.1 (R Core Team (2017).

## Results

### Assay development and *in-silico* and *in-vitro* validation

Primers and probe were highly species-specific as *in-silico* and *in-vitro* tests did not reveal any matches with non-target species (Table S3). Analysis of the standard curve (Fig. 1A) revealed a strong dependency of Ct values on DNA concentrations (y=-1.73x+20.8, *p*<0.001, *r*^2^ = 0.996). Likewise, the detection probability was also positively related to DNA concentration in the sample (y=-0.18x+1.39, *p* = 0.0016, *r*^2^ = 0.804; Fig. 1B), highlighting the possibility of a quantifiable assay being developed. Method sensitivity analyses revealed a LOD of 7.9 x 10^-5^ng and a LOQ of 7.9 x 10^-4^ng crayfish DNA extract per μL^-1^.

**Figure 1.**
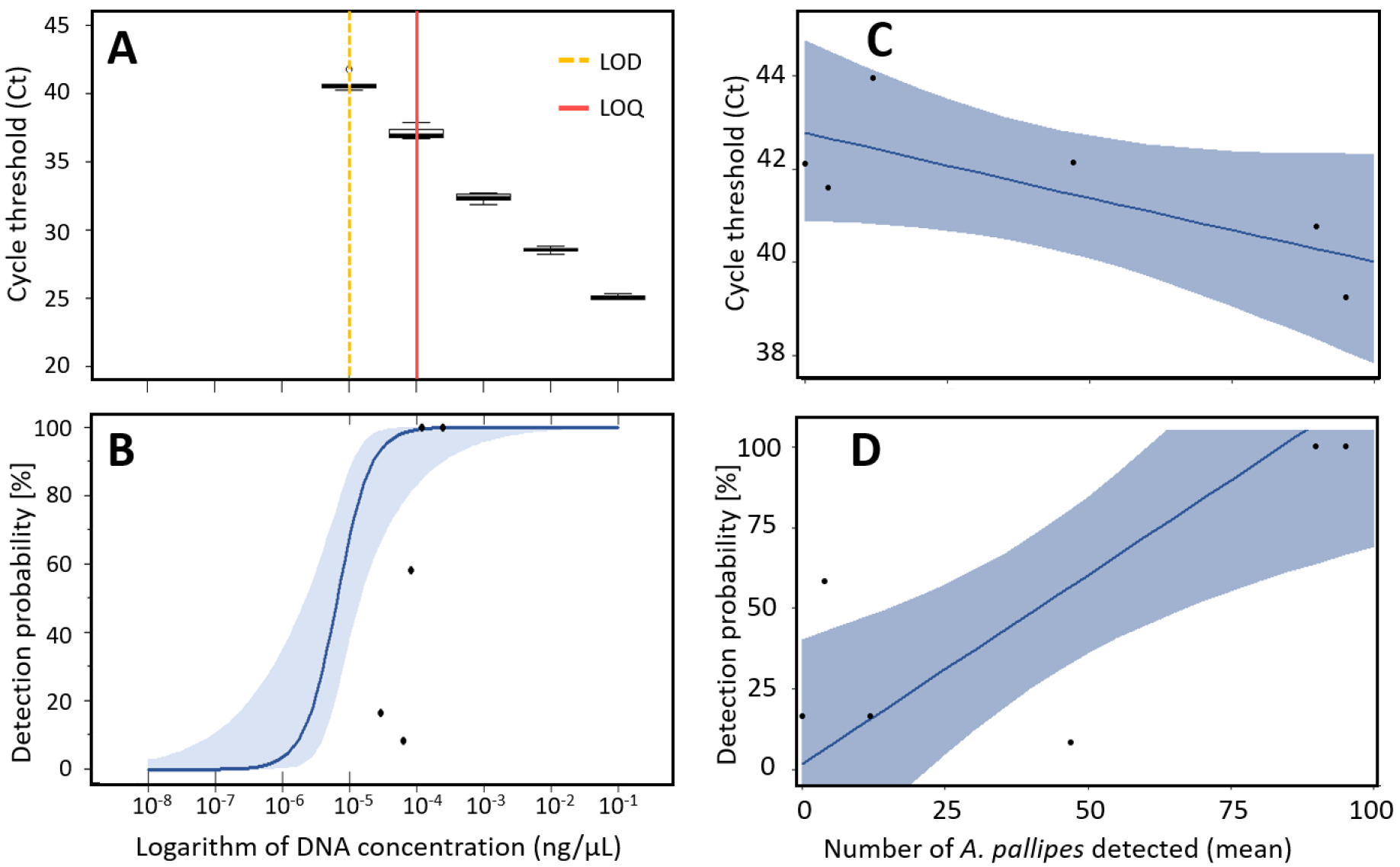
**(A)** Relationship between cycle threshold (Ct) and DNA concentration from White-clawed crayfish qPCR calibration curve. Limit of detection (LOD) and limit of quantification (LOQ) are illustrated by vertical lines (dashed-yellow and red respectively). **(B)** Change in detection probability with increasing DNA concentrations in calibration curve data (blue line). Red triangles represent field data (in-*situ* validation data) that is compared with the log-regression based on in-*vitro* calibration curve data. Three out of five field data points were outside the established confidence interval, indicating discrepancies between field and laboratory-based data sets. **(C)** Relationship between Ct values and white-clawed crayfish population monitored using traditional method. **(D)** Relationship between detection probability of eDNA and traditionally evaluated crayfish population sizes. The blue line and the light-blue shaded area reflect the results of a logit regression and its 95% confidence interval, respectively.

### In-situ validation

Populations of white-clawed crayfish were found in five out of the six surveyed sites using traditional survey methods. eDNA-based detection indicates the presence of white-clawed crayfish in all six sites, though the site with no visual white-clawed crayfish sightings was characterised by a very low detection probability. The Ct values from the six river sites were converted into DNA concentrations using the calibration curve, which allowed us to compare the relationship between detection probability and DNA concentration in laboratory and field samples (Fig. 1B). Four out of the six field sites lay outside of the 95% confidence interval of the standard curve, indicating systematic differences between *in-vitro* validation and field samples. As a direct result of limited recapture data, no recapture was observed in three out of the six survey locations despite indication of the presence of a significantly sized population. Estimates of population sites were therefore unable to be reliably made using the Lincoln-Petersen approach. The relationship between the mean number of crayfish detected across all sampling sates was therefore applied instead. The relationship of the mean number of crayfish detecting using conventional species survey methods (torching) and detection probability of eDNA measurements (Fig. 1D) was significant, but only when water temperature was included (y=0.0118x_1_-0.117x_2_+1.77; x_1_=mean survey count, x_2_=temperature, *p*=0.035, *r*^2^=0.82, best model resulted also in reduction of AIC by 10 units), with two out of the six datapoints lying outside of the confidence interval. The relationship between Ct and the mean number of crayfish detected using torching was marginally non-significant but showed a reasonable model fit (Fig. 1C; y=-0.00067log(x)+3.76, *p*=0.079, *r*^2^=0.47, best model resulted in reduction of AIC by 5 units). Differences in filtered sample volume did not significantly influence results.

### Comparison of eDNA sampling methods

In mesocosm experiments, sampling methodology had a significant impact on detection probability (ANOVA F_(3,44)_=74.48, *p*<0.001). Pairwise comparisons revealed that detection probabilities of all three filtration-based methods (2μm, 0.22μm and 0.45μm) were comparable (*p*>0.05) but differed significantly from the precipitation method (*p*<0.001, Fig. 2A). However, the *p*-value for the comparison between 0.45μm and 2μm was marginally non-significant (*p*=0.051). Similarly, methodologies also differed significantly in Ct (ANOVA F_(3,178)_=90.1, *p*<0.001). However, in contrast to detection probability, pairwise tests indicated a difference between the 2μm filtration method and all the other approaches (p<0.001; Fig. 2B; only samples with positive detection were included in the analysis).

**Figure 2.**
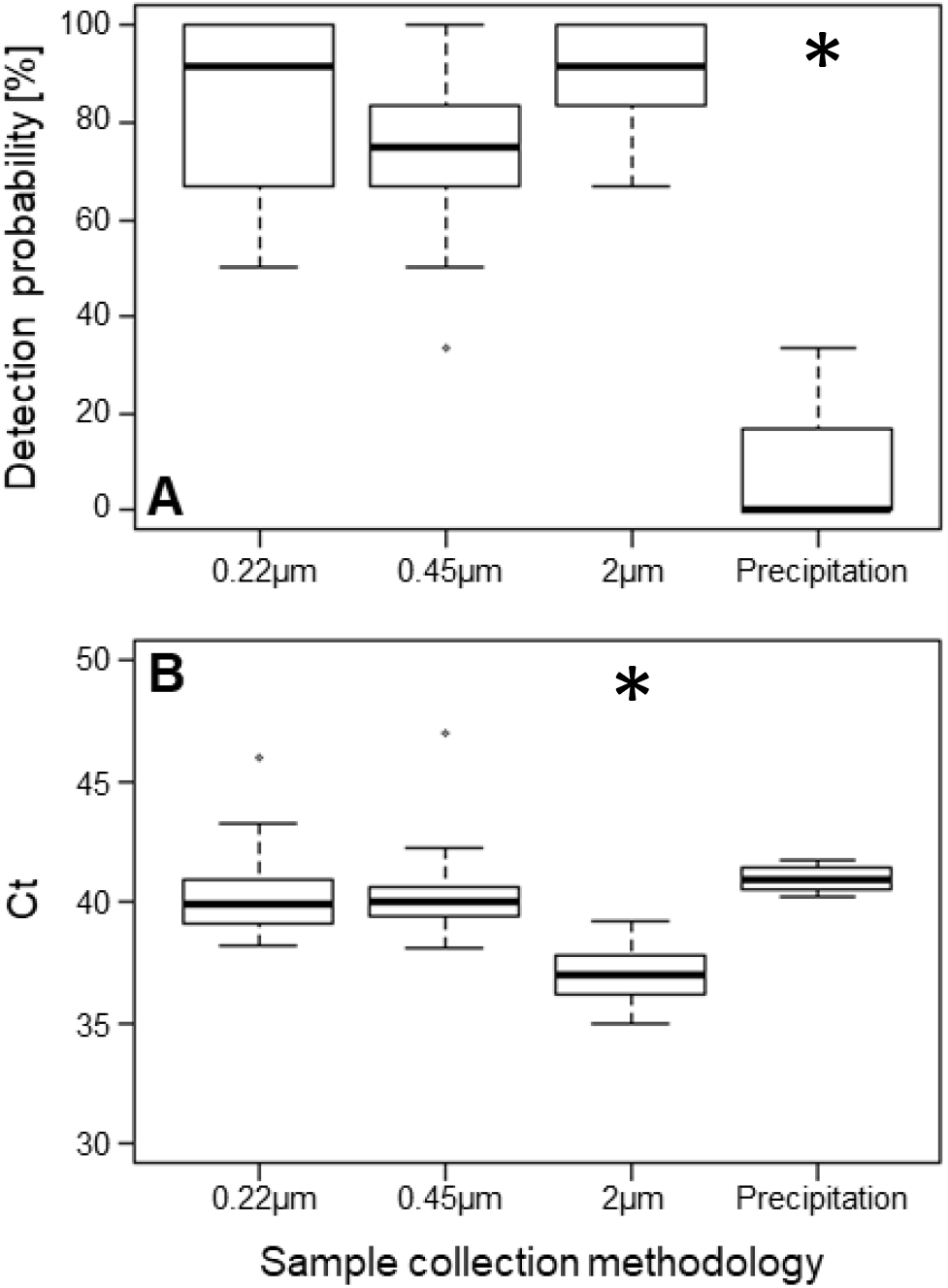
Comparison of the detection probability **(A)** and Ct values **(B)** of white-clawed crayfish using different eDNA sampling methods (0.22μm filtration, 0.45μm filtration, 2μm filtration and precipitation) in a controlled mesocosm experiment (* indicates statistical significance).

*In-situ* comparisons of sampling methods in a lentic system were highly comparable to the mesocosm experiment (Fig. 3 A-B). The precipitation method showed a significantly lower detection probability (T-test, t=3.55, df=75.37, *p*<0.001) and a significantly higher Ct (t=-2.46, df=15.72, *p*<0.05) than the filtration-based method (0.22μm). However, contrasting results were attained in lotic systems. Here, we assessed the method for both, white-clawed crayfish and the crayfish plague (not present in mesocosms or ponds). The detection probability of crayfish plague mirrored findings from other systems showing significantly higher detection probabilities for the 2μm filtration method (nested ANOVA; F_(1,69)_=4.92, p<0.05; Fig. 3E). Ct values were not significantly different, but also indicated a better performance of the filtration-based method (Fig. 3F). However, the results for white-clawed crayfish contrasted all other results. In lentic systems, precipitation resulted in a higher detection probability (nested ANOVA F_(1,69)_=13.77, *p*<0.001, Fig. 3C) and accordingly, lower Ct values (nested ANOVA; F_(1,34)_=5.24, *p*=0.028; Fig. 3D). Consequently, filtration-based methods performed consistently better except in lentic systems where eDNA from white-clawed crayfish was more reliably assessed with the precipitation method.

**Figure 3.**
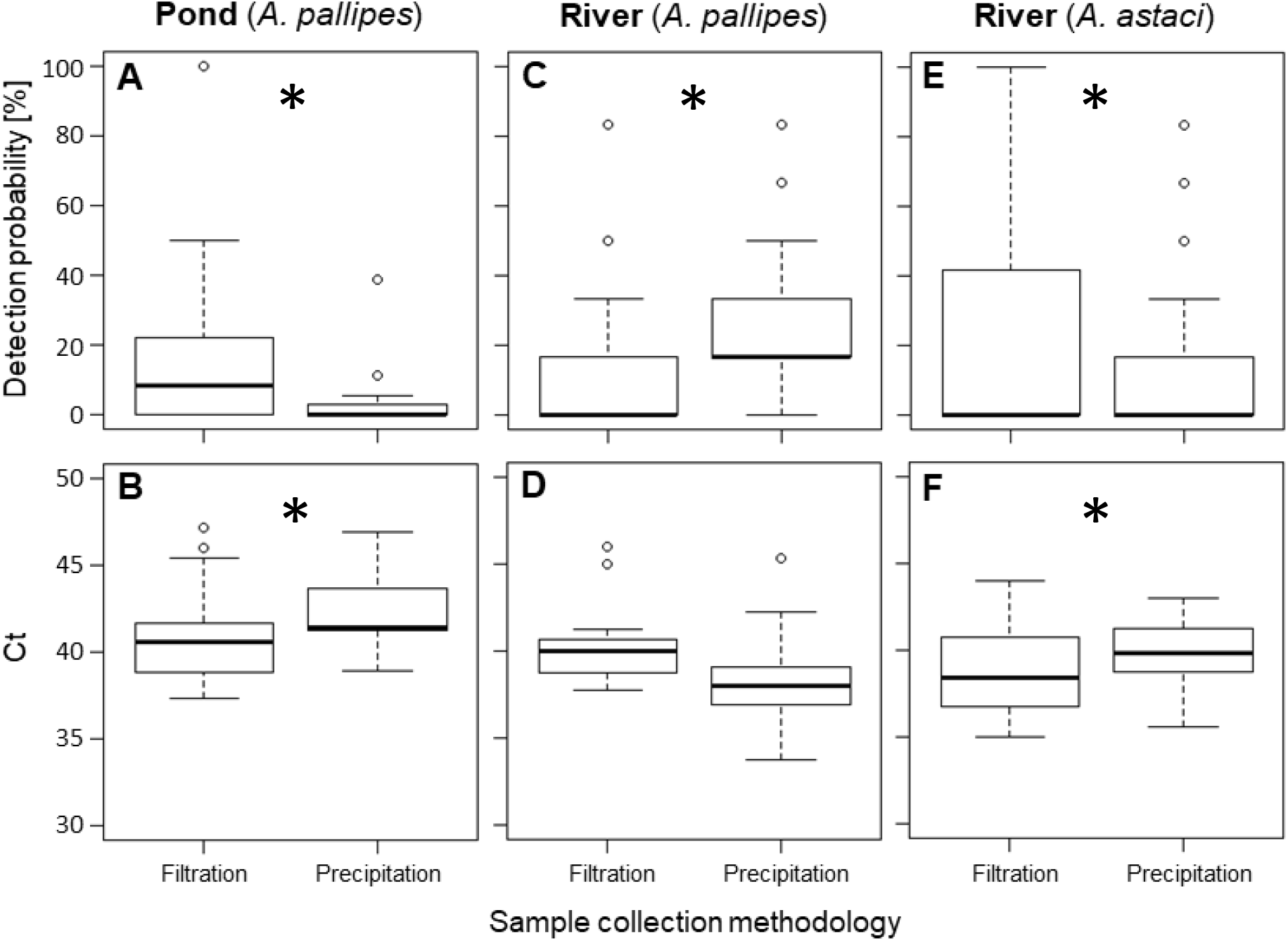
Comparison of the detection probability **(A, C, E)** and Ct values **(B, D, F)** of different eDNA sampling methods (filtration and precipitation) for white-clawed crayfish in a lentic system **(Pond, A-B)** (filter pore size 0.22 μm) and for both white-clawed crayfish **(River, C-D)** and crayfish plague **(River, E-F)** in the same lotic system (filter pore size 2 μm) (* in panels signifies significant differences between pairwise method).

### Field tests of species co-occurrence

Finally, our joint assessment of white-clawed crayfish, signal crayfish and crayfish plague (Fig. 4) demonstrated that white-clawed crayfish occurrence was related to both other species (Fig. 4, B,C). Whilst univariate regressions were marginally non-significant (dependency of white-clawed crayfish on signal crayfish: *p*=0.063; dependency of white-clawed crayfish on crayfish plague: *p*= 0.051), a multiple regression analysis revealed significant relationships (y = −23.8x_1_ + 13.1x_2_ - 3.8, *r*^2^=0.73, *p*=0.016; y, x_1_ and x_2_ represent detection probabilities of white-clawed crayfish, signal crayfish and crayfish plague, respectively). Unsurprisingly, white-clawed crayfish was negatively impacted by the presence of signal crayfish (Fig. 4B), yet contrary to expectation they were shown to be positively related with increase in detection of crayfish plague (Fig. 4C). There was no apparent correlation between signal crayfish and plague (Fig. 4D).

**Figure 4.**
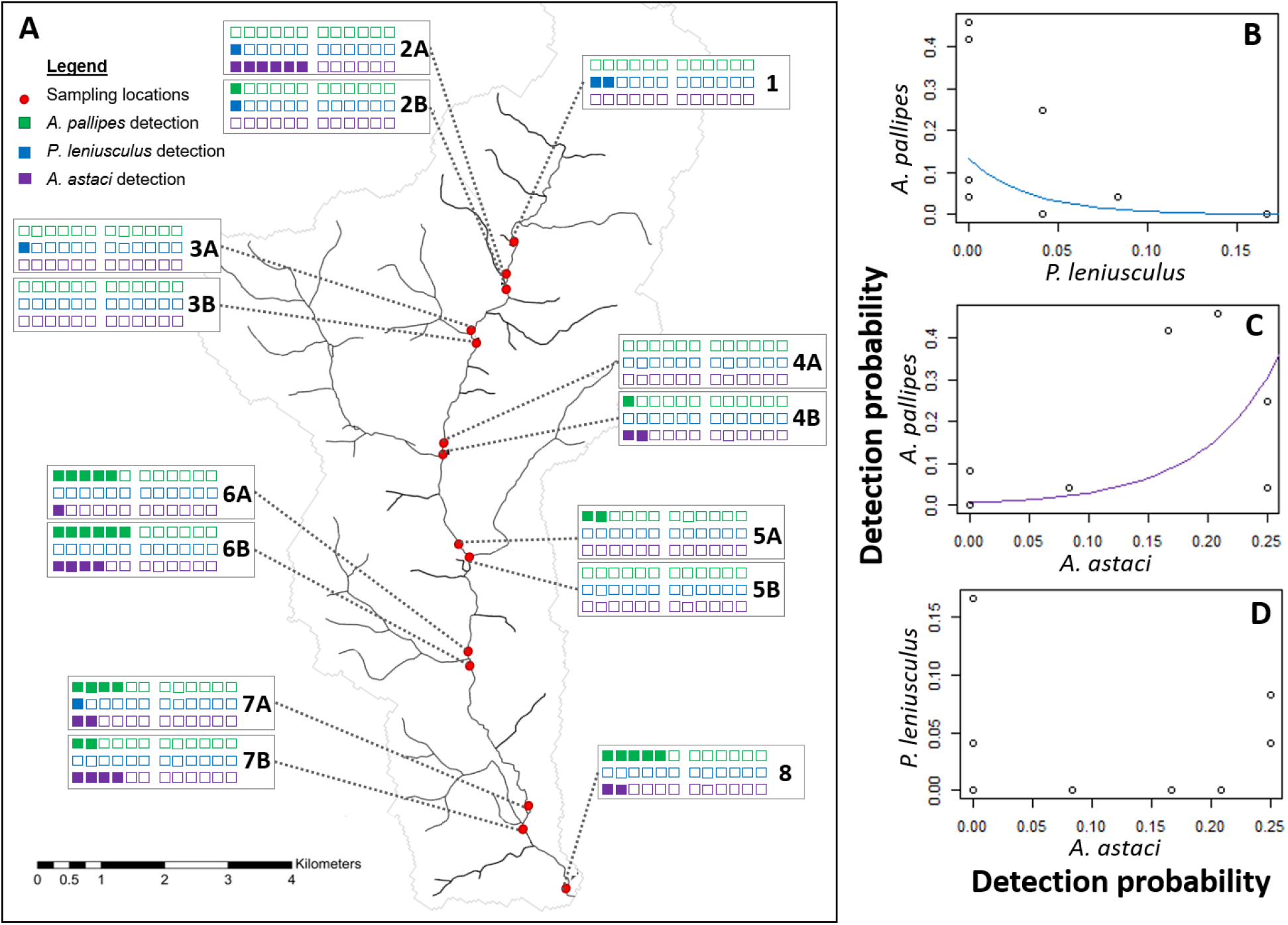
**(A)** Detection of eDNA from white-clawed crayfish (green squares), signal crayfish (blue squares) and crayfish plague. (purple squares) in a river catchment in Derbyshire. Eight locations were sampled and are represented by red dots. The empty squares represent the negative qPCR replicates. **(B)** Indicates the relationship between the detection probability of white-clawed crayfish and detection probability of signal crayfish. **(C)** The relationship between the detection probability of white-clawed crayfish and detection probability of crayfish plague. **(D)** The relationship between the detection probability of signal crayfish and detection probability of crayfish plague.

## Discussion

Native crayfish species across Europe are threatened by invasive competitors and the crayfish plague, resulting in a downward trajectory of native species’ abundance and distribution (Holdich et al., 2009). In this study, we present a novel assay for the detection of white-clawed crayfish, a flagship conservation species in Western Europe. In rigorous *in-vitro* and *in-situ* tests, we evaluated the reliability of our assay under various environmental conditions. Further, we applied our assay together with established eDNA-based methods to assess the drivers of white-clawed crayfish occurrence. Overall, we were able to demonstrate that our approach can not only be used for simple presence/absence surveys but also has the potential to reveal species co-occurrence and interactions. However, our results also highlight that such applications are only meaningful after thorough method testing and validation.

Field comparisons indicated a higher sensitivity of the eDNA assay compared to conventional surveys, which only resulted in positive detection in five out of six sites. Such traditional surveys are reported to vary in detection success (Gladman et al., 2010), therefore making comparisons between eDNA and these methods challenging. Additionally, the two UK lotic river systems used to compare different sampling approaches were since conventionally surveyed for the presence of crayfish, with no white-clawed crayfish found, despite previous recorded presence, suggesting a higher sensitivity of the eDNA assay. Whilst higher sensitivity is frequently reported for eDNA assays (Dejean et al., 2012; Smart et al., 2015), and therefore likely represent a more reliable species detection approach, such results should be interpreted with caution as eDNA-based approaches are associated with a risk of providing false positive results (Furlan et al., 2016), for example via downstream transport of eDNA within river networks (Pont et al., 2018). Moreover, false positives may result from historic eDNA, which is still present after the extinction or emigration of the target species (Turner et al., 2015). In our case, this represents a valid hypothesis as all field sites were populated by white-clawed crayfish a year before our field surveys (C. Mauvisseau, personal communication). Consequently, it remains inconclusive whether the developed eDNA approach truly has a higher sensitivity (i.e. false negative of torching method) or, in fact, white-clawed crayfish was not present at the field site in question.

Further, an important component of our method validation was the comparison of different field sampling approaches. Precipitation and filtration protocols to concentrate eDNA from the environment have already been compared in a number of studies (Deiner et al., 2015; Spens et al., 2017). Most investigators endorse filtration approaches (Hinlo et al., 2017b; Spens et al., 2017; Vörös et al., 2017), but optimal pore size may differ between species (Spens et al., 2017). Moreover, method choice can also be environment-dependant. For example, Eichmiller, et al. (2016) indicated filtration as the optimal eDNA-based method for surveying the common carp, *Cyprinus carpio*. In contrast, Minamoto et al. (2016) highlighted precipitation performed better – a result likely brought about by variations in the environment across both studies. In our controlled mesocosm comparison, we found that a 2 μm filtration approach outperformed precipitation and the other filtration methods tested. However, field comparisons revealed contrasting results, again likely brought about by the different environments surveyed (Fig. 3). In this scenario precipitation outperformed filtration (2 μm) in the lotic system.

One possible explanation for our divergent findings across different habitats is that target eDNA particles differ in these environments. eDNA is exposed to continuous degradation through biotic (e.g. bacteria) and abiotic (e.g. UV) factors (Strickler et al., 2015) and these degradation processes can affect eDNA particle size distributions. Filtration has the advantage to collate eDNA from larger water volumes but is linked to the risk of losing particles which are below filter pore sizes. Hence, the habitat-specific differences in our method comparisons may be explained by the specific degradation processes within the investigated river systems. A decrease of average eDNA particle size below the filter pore size would substantially decrease detection probability of filtration approaches and explain our findings.

An alternative explanation for our results is linked to inhibition of eDNA amplification. Inhibitor compounds (that interfere with qPCR processes), have been shown to affect target DNA amplification in a non-linear way (Goldberg et al., 2016). If inhibitor concentration is low, amplification will not be strongly impacted. However, if concentrations surpass a certain threshold, inhibitors may suppress the amplification of even high concentrations of target eDNA (Mauvisseau et al., 2019a). Sampling methods that differ in their water collection volumes and in the amount of concentrated target eDNA, will also concentrate inhibitors to different degrees (Fig. 5). Consequently, sampling methods that reach higher target eDNA concentrations may show a lower overall performance due to the non-linear relationship between inhibitor concentrations and DNA amplification. This scenario will occur when inhibitors are present in high concentrations and efficiently concentrated. Therefore, different ratios between target eDNA and inhibitors in different environments can cause a shift in the relative performance of sampling methods across habitats (Fig. 5). In our case, we did not include tests for inhibition, which include the addition of synthetic DNA to qPCR reactions (i.e. failure to detect synthetic DNA indicates inhibition; (Goldberg et al., 2016; Mauvisseau et al., 2019a). However, we recommend that such inhibition tests should be included in future field method comparisons.

**Figure 5.**
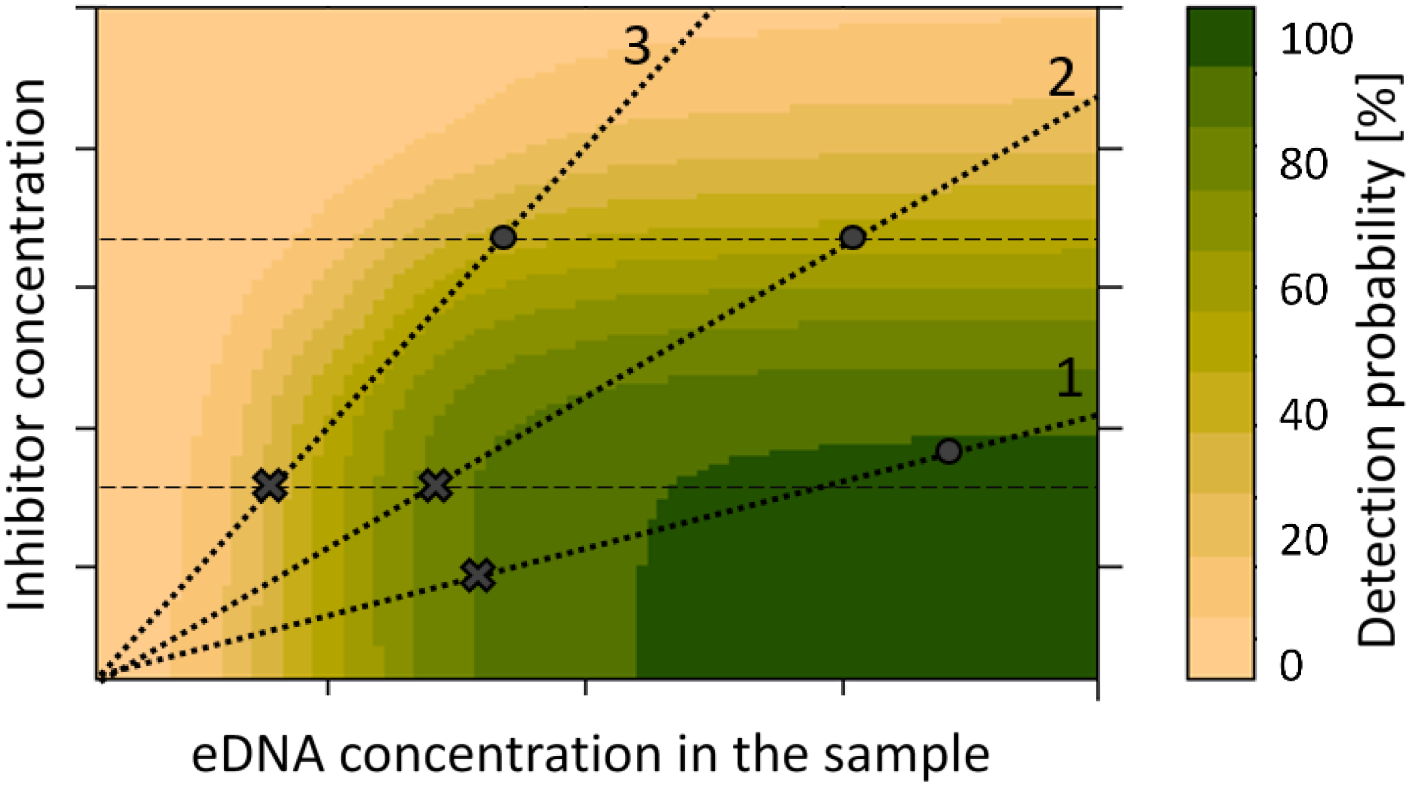
Schematic of the co-dependency of detection probability on target eDNA and inhibitors concentrations in water samples. Detection probability increases with eDNA concentration and decreases with inhibitor concentrations but is low when both variables are high. Each water body is characterised by a certain ratio between inhibitor and target eDNA concentrations represented by black dotted lines (**1-3**). A change in sampling methods accompanied by a change in the sampled water volume will result in different concentrations of target eDNA and inhibitors in the sample and in shifts along dotted lines (**grey crosses and dots**). An increase in sampled water volume will therefore in some water bodies increase (Line 1) and in other decrease (Line 2) detection probability. The same is true when different eDNA assays in the same water body are considered. While eDNA concentrations of two targets may differ, inhibitor concentrations will be the same. Consequently, samples with the same water volume will have the same inhibitor concentrations (horizontal dashed lines). Nevertheless, changes in sampling volume and method can result in increased detection probability for one target (Line 3) but not for the other (Line 2).

Both inhibition and different target-eDNA size distributions might also explain differences in method comparisons between species in the same environment as observed for white-clawed crayfish and crayfish plague in lotic habitats (Fig. 3). A fundamental distinction between the two species is that crayfish plague depends for its proliferation on the frequent and abundant release of encapsulated spores (~8 μm in diameter). It seems likely that these spores, which are designed for transport along large distances, will show lower sensitivity to degradation than white-clawed crayfish DNA, which potentially could explain our species-specific results.

Finally, we demonstrated that our approach can also be used for investigating ecological relationships determining the distribution of endangered species. Our simultaneous assessments of white-clawed crayfish, signal crayfish and crayfish plague revealed a negative impact of signal crayfish on white-clawed crayfish (Fig. 4). Such negative impacts of invasive competitors on native crayfish species have been frequently highlighted before (Holdich et al., 2009) and demonstrate the applicability of our approach. Interestingly, however, we illustrated a positive relationship between white-clawed crayfish and crayfish plague, which went against our expectation. Such co-occurrence might result from the one-time nature of our sampling approach and reflect a disease outbreak within the crayfish population, which most probably would result in local extinction (Strand et al., 2019). However, recent discoveries have also indicated the potential of plague resistance in some white-clawed crayfish strains (Martín-Torrijos et al., 2017). Such increased disease tolerance might facilitate a permanent co-existence of pathogen and host. Consequently, further in-depth monitoring of species dynamics together with genetic profiling and disease susceptibility tests should be a primary objective of future conservation planning.

### Conclusions

Currently, many species-specific eDNA assays only cover *in-silico*, *in-vitro* and sometimes basic *in-situ* validation steps (Dickie et al., 2018; Lacoursière-Roussel et al., 2016). Already published white-clawed crayfish eDNA assays have shown some promising first results but yet need to go through the required thorough level of *in-situ* evaluation (Atkinson et al., 2019; Robinson et al., 2018). Here we illustrate that sampling methods can differ strongly in performance and recommend rigorous testing of eDNA assays to optimise sampling strategies. However, our contrasting results of method comparisons across habitats and species highlight that there might not be something like a universal ‘optimal sampling method’, but that adjustments to account for local conditions are required. The resulting higher method reliability increases the applicability of eDNA assays and paves the way for more detailed ecological studies to improve species management and conservation.

## Acknowledgments

For assistance with sample collection we thank Holly Thompson, Jacob Ball, Jack Greenhalgh, Isabelle Parot and Antoine Monier. We would also like to thank Alex Lumsdon and the Environment Agency, Angus Menzies and Dorset Wildlife Trust for funding and their assistance with sample collection. For providing DNA samples: Catherine Souty-Grosset, Sina Tönges, Frank Lyko, Jenny Makkonen and Japo Jussila. Funding was provided by SureScreen Scientifics, UK.

## Author Contributions

C.T., M.S. and J.N. designed the experiment and methodology, C.T., Q.M., J.N. and C.M. collected field samples, C.T., and Q.M. performed extraction and PCR, C.T., A.B. and M.B. analysed the data. The manuscript was written by C.T., M.S. and A.B. and reviewed by all authors.

